# A cell membrane-hybridized nanocarrier-based RNA delivery system for ocular neovascular diseases

**DOI:** 10.64898/2025.12.23.696136

**Authors:** Bo-Jiao Tang, Man-Hong Li, Xian-Chun Yan, Yu‑Sheng Wang, Jia‑Xing Sun

## Abstract

Ocular neovascularization is a common pathological feature of various blinding eye diseases. Current anti vascular endothelial growth factor (VEGF) therapies face challenges such as low response rates, frequent injections, and multiple complications. RNA-based therapeutics offer advantages including strong targeting capability and reversible modulation, but encounter obstacles like low delivery efficiency, susceptibility to degradation, and lack of targeting specificity. This study developed an endothelial cell membrane-hybridized lipid nanoparticle that achieve specific targeting of neovascular endothelial cells through “homing effects”. The carrier exhibits excellent biocompatibility, high small interfering RNA (siRNA) delivery efficiency, and lysosomal escape capability, effectively penetrating the vitreous barrier. In a laser-induced choroidal neovascularization mouse model, it significantly enhanced drug accumulation at the lesion site, providing a safe and effective delivery strategy for RNA therapy of ocular neovascular diseases.

## Introduction

Ocular neovascularization is a common pathological feature of various refractory blinding eye diseases, including diabetic retinopathy (DR) and age-related macular degeneration (AMD), leading to visual impairment and irreversible blindness ^1^. With the increasing incidence of diabetes, population aging, and other lifestyle factors, the prevalence of ocular neovascularization has risen significantly, exerting a major impact on global visual health ^2^. Though anti-VEGF therapy serves as the first-line clinical treatment for inhibiting neovascularization, over 60% of patients still exhibit disease progression, manifesting as poor response to therapy ^3^. In addition, the requirement for frequent injections not only lead to poor patient compliance but also pose problems such as devastating ocular complications (e.g., endophthalmitis, intraocular hemorrhage, retinal tears, and retinal detachment) ^4^. Therefore, exploring novel therapeutic strategies to more effectively control ocular neovascularization and improve patient prognosis has become a key focus of current research.

Compared to traditional protein-targeting and DNA-based drugs, RNA-based therapeutics demonstrate promising potential due to their unique physicochemical and physiological properties^5^. RNA plays functional roles among all three major biomolecules (DNA, RNA, and proteins)^5^. Therefore, theoretically, RNA-based drugs can be designed to target any gene of interest by selecting a complementary nucleotide sequence ^6^. RNA therapy functions by targeting RNA or proteins, encoding missing or defective proteins. It enables specific and reversible genetic manipulation independent of DNA, thereby avoiding permanent alterations in the host organism ^7^. Therefore, targeting messenger RNA (mRNA) significantly broadens the treatment.

Despite the broad prospects, delivery of mRNA face significant challenges. Their large size, negative charge, and susceptibility to ribonuclease in blood and tissues hinder their efficient cellular uptake and intrinsic activity ^8^. Moreover, the vitreous humor is characterized by high water content (>98%) and contains a negatively charged macromolecular matrix composed of collagen fibers, hyaluronic acid, and glycosaminoglycans. Most substances injected into the vitreous cavity tend to be retained within this matrix, making it difficult to penetrate the retina ^9^. Moreover, due to potential off-target effects that may impact other critical pathways, RNA therapeutics rely on delivery systems to achieve targeting specificity and enhance therapeutic efficacy ^10^. However, previous viral vector-based delivery systems, including adeno-associated viruses, adenoviruses, and lentiviruses, all have limitations such as safety concerns, limited gene cargo capacity, and high costs ^11^.

The nanoparticle drug delivery systems mainly include lipid-based nanoparticles and polymer nanoparticles ^8^. The main advantage of lipid-based nanoparticles lies in their customizable optimization of biophysical properties (such as size, shape, and chemical/material composition) and biological properties (such as targeting ligand functionalization), thereby enabling the construction of highly specific delivery platforms ^12^. Based on this, the present study constructed an endothelial cell membrane-hybridized lipid nanoparticle (Cell membrane-Lips) for siRNA delivery. This nanoparticle exhibits low surface positive charge and excellent lysosomal escape properties, making it suitable for traversing the vitreous cavity to reach the retina. By leveraging the “homing effect” of cells, it targets neovascular lesions to achieve endothelial cell-specific, efficient, and safe silencing of the target gene. Using a laser-induced choroidal neovascularization (CNV) mouse model and in vitro experiments, we demonstrated that intravitreal injection of Cell membrane-Lips can effectively deliver siRNA to neovascular lesions, providing a novel strategy for drug development in intraocular neovascular diseases.

### Experimental Section

#### 1. Materials

The polycarbonate membrane was purchased from Whatman (Britain). Soybean phosphatidylcholine (SPC), cholesterol, and ((4-Hydroxybutyl) azanediyl) bis (hexane-6,1-diyl) bis (2-hexyldecanoate) (ALC-0315) were provided by Ruixi (China). The siRNA (si NC-cy3) was purchased from Gene Pharma (China). Anti-adhesion G protein-coupled receptor E1 (Adgre1, also known as F4/80) was purchased from Cell Signaling Technology (USA), anti-solute carrier family 1 member 3 (Slc1a3, also known as GLAST) was purchased from Proteintech (China), and anti-cadherin 5 (Cdh5, also known as VE-cadherin) antibody was purchased from Abcam (UK). Griffonia Simplicifolia Lectin I isolectin B4 (IB4) (FL-1201, Fluorescein) was purchased from vector laboratories (USA). Redi Plate™ 96 Ribo Green® RNA Quantitation Kit (R-32700), Lyso Tracker™ Green DND-26 (L7526), and donkey anti-goat IgG H&L (Alexa Fluor 488) (A-11055) were purchased from Invitrogen. DAPI was purchased from Beyotime (China). Ava was provided by Roche. Cell Counting Kit-8 (CCK-8) reagents were obtained from YEASEN (China). Cell lines were provided by ATCC. Mammalian cell culture medium, fetal bovine serum (FBS), optimal cutting temperature (OCT) compound, and 4% paraformaldehyde (PFA) were purchased from Abcam (UK).

#### 2. Synthesis and Characterization of Cell membrane-Lips

##### 2.1 Synthesis of Cell membrane-Lips

Dissolve SPC, ALC-0315, and cholesterol together in 3 mL of anhydrous ethanol, then transfer to an eggplant-shaped flask. Dissolve si NC-cy3 in a citric acid buffer (50 mM citrate, pH 4) containing 25% ethanol and slowly add it to the lipid mixture with thorough mixing, followed by incubation for 20 minutes. Extraction of cell membranes through sonication and liposome extruder (100 nm filter membrane) treatment. The prepared liposomes were mixed with macrophage cell membranes, Müller cell membranes, and endothelial cell membranes at a mass ratio of 1:1, respectively, followed by water bath sonication to promote fusion between the liposomes and cell membranes until the solution became transparent. This resulted in the synthesis of macrophage membrane-hybridized liposomes (Macrophage-Lip), Müller membrane-hybridized liposomes (Müller-Lip), and endothelial membrane-hybridized liposomes (Endothelium-Lip). Lyoprotectants were then added for freeze-drying.

##### 2.2 Encapsulation efficiency and drug - loading capacity

The encapsulation efficiency (EE%) and the drug - loading capacity (LC (wt%)) were calculated according to Equation 14 reported previously. Formulas

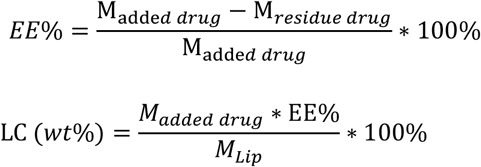

M_added drug_ and M_residue drug_ represent the original mass of siRNA and the mass in the supernatant, respectively; M_Lip_ is the weight of the synthesized liposomes. The mass of siRNA was determined using the Redi Plate™96 Ribo Green® RNA Quantitation Kit and repeated three - times.

##### 2.3 Characterization

The particle size and zeta potential measurements were performed by diluting the Cell membrane-Lips and taking 1 mL to add into the particle size cell, using the Nano Brook 90plus PALS nanoparticle size and zeta potential analyzer (Brookhaven Instruments, USA). The morphology of the liposomes was observed using transmission electron microscopy (TEM, Talos F200S FEI).

##### 2.4 Identification of Cell-Membrane Components

Cell membrane-Lips were mixed with Loading buffer (Beyotime), incubated at 95°C in a metal bath for 5 minutes, and then separated by sodium dodecyl sulfate-polyacrylamide gel electrophoresis (SDS-PAGE). Immunoblotting was performed using primary antibodies including anti-F4/80 antibody, anti-GLAST antibody, and anti-VE-cadherin antibody incubated on polyvinylidene fluoride (PVDF) membranes, followed by incubation with horseradish peroxidase (HRP)-conjugated secondary antibody (1:2000). After each step, wash the membrane three times with TBST containing 0.1% Tween-20 for 10 minutes each. The membrane was visualized using an enhanced chemiluminescence system (Clinx Science Instruments, China).

#### 3. Cytotoxicity Test

Mouse brain microvascular endothelial cells (bEnd.3) were seeded in 96-well plates at a density of 5×10^3^ cells/well and cultured in a 37°C, 5% CO2 incubator for 24 hours. After discarding the culture medium, the cells were washed three times with sterile 1×PBS. Each well was supplemented with media containing different concentrations of Macrophage-Lip, Müller-Lip, and Endothelium-Lip (0, 150, 250, 500, 1000 μg/mL), followed by continued incubation in the cell culture incubator for 24 h. Remove the cell culture plate, discard the medium, wash the cells three times with sterile 1×PBS, add medium containing 10% CCK-8 reagent, continue to incubate in the cell culture incubator protected from light for 2 hours, then measure the absorbance at 450 nm using a microplate reader. Set up duplicate wells for each group, observe and calculate cell viability.

#### 4. Cellular Uptake

bEnd.3 cells were seeded in 24-well plates at a density of 5×10^4^ cells/well and cultured in a 37°C, 5% CO_2_ incubator for 24 hours. After complete cell adhesion, the cells were washed with sterile 1×PBS. Subsequently, medium containing Macrophage-Lip, Müller-Lip, and Endothelium-Lip (each at 150 μg/mL) was added separately, followed by continued culture for 24 hours. The medium was then discarded, and the cells were fixed with 4% PFA for 20 minutes, followed by 3 times of PBS washes. Hoechst was added for 10 minutes of staining, and then anti-fade mounting medium was applied for sealing. The prepared samples were observed and photographed using confocal microscopy (FV 3000, Olympus, Japan).

#### 5. Lysosomal Escape

bEnd.3 cells were seeded in a 24-well plate at a density of 5×10^4^ cells per well and cultured for 24 hours. Medium containing Endothelium-Lip (150 μg/mL) was added, and the cells were cultured for an additional 2 hours or 8 hours, respectively. After incubation, the cells were washed with PBS, followed by the addition of LysoTracker diluted in DMEM medium to each well. The plate was then incubated in the dark for 2 hours in a cell culture incubator. Subsequently, the cells were washed again with PBS and stained with Hoechst for 10 minutes, and then mounted using anti-fade mounting medium. Lysosomal escape was observed and photographed using confocal microscopy (FV 3000, Olympus, Japan). Colocalization analysis was performed with Image J to calculate Manders’ Colocalization Coefficients (MCC).

#### 6. *In-vivo* Experiments

##### 6.1 Animal

Six-to-eight-week-old male C57BL/6J mice (SPF grade) with average body weight of 18-22 g and good general condition were used. The mice were housed under a 12 hours / 12 hours light-dark cycle and fed standard diet with free access to food and water during the experiment. All animal procedures followed the Association for Research in Vision and Ophthalmology (ARVO) Statement for the Use of Animals in Ophthalmic and Vision Research. Prior to experiments, animal eyes were examined using slit-lamp microscopy to exclude anterior segment abnormalities. The study protocol was approved by the Animal Welfare and Ethics Committee of The Fourth Military Medical University (Approval No. 20250003).

##### *6*.*2 In-vivo* biocompatibility experiments

Prior to conducting animal experiments, the intraocular biocompatibility of Cell membrane-Lips in C57BL/6J mice were studied. Twelve mice were randomly divided into 4 groups. Each vitreous cavity was injected with 1 μL of Macrophage-Lip, Müller-Lip, and Endothelium-Lip. The control group received 1 μL of normal saline. On day 7, the eyeballs were enucleated, embedded in paraffin, and retinal histological changes were observed using hematoxylin-eosin (HE) staining.

##### 6.3 Establishment of the CNV model in mice

Sixteen C57BL/6J mice were randomly divided into 4 groups: si NC-cy3 group, Macrophage-Lip group, Müller-Lip group, and Endothelium-Lip group. After one week of acclimatization, mice were anesthetized via intraperitoneal injection of 1% sodium pentobarbital solution. Pupils were dilated with compound tropicamide eye drops, and a corneal contact lens was placed. Using a slit-lamp microscope with a 532-nm frequency-doubled laser (spot diameter: 50 μm, power: 90 nW, exposure time: 100 mS), five uniform photocoagulation spots were applied per eye around the optic nerve. Successful Bruch’s membrane rupture was confirmed by bubble formation or mild sound, which served as the criterion for valid laser spots.

##### 6.4. Immunofluorescence staining

On day 7, choroidal flat mounts were prepared for CNV volume analysis. The choroid-sclera complex tissues were post-fixed in 4% PFA at 4°C for 6 hours, then blocked in PBS containing 1% bovine serum albumin (BSA) and 0.5% Triton X-100 at room temperature for 2 hours. Subsequently, the tissues were incubated with IB4 antibody in blocking buffer at 4°C overnight, followed by incubation with secondary antibodies in PBS at room temperature for 2 hours. After each step, the samples were washed 3 times with PBS. The choroidal was flat-mounted on glass slides and cover slipped with 50% glycerol. Imaging was performed using confocal microscopy (FV 3000, Olympus, Japan).

#### 7. Statistical Analysis

Statistical analysis was performed using Image-Pro Plus6.0 and GraphPad Prism 8.0. All quantitative data are presented as means ± SD. Statistical significance was calculated using Student’s t-tests for continuous variables between two groups. Comparison of continuous variables among more than two groups was performed by one-way ANOVA followed by Tukey’s post hoc test for one independent variable. Nonparametric tests were used for non-normally distributed data. Probability values were two-tailed and p < 0.05 was considered statistically significant.

## Results

### 1. Synthesis and Characterization of Cell membrane-Lips

Ocular neovascular lesions consist of various cellular components such as endothelial cells, macrophages, and Müller cells ^13 14^. Therefore, we prepared three types of hybrid liposomes - namely, Macrophage-Lip, Müller -Lip, and Endothelium-Lip - and compared their targeting efficacy toward neovascular lesions. The particle size and zeta potential of the three liposomes were measured immediately. As shown in Figure 1 (a-c), all three liposomes were in the nanometer range with small polydispersity index values and uniform dispersion, which facilitates homogeneous distribution after intravitreal injection;Among them, Macrophage-Lip had the smallest average particle size of approximately 106 nm, while Müller -Lip measured 132 nm and Endothelium-Lip was 124 nm in size. The potential diagram as shown in Figure 1(d) indicates positive potential, all around +10mV. Since the vitreous cavity contains negatively charged substances such as collagen fibers, hyaluronic acid, and glycosaminoglycans ^13^, coating the liposome surface with cell membrane helps mitigate their excessively strong positive charge, thereby enhancing their efficiency in traversing the vitreous cavity and reaching the retina.

**Figure 1.**
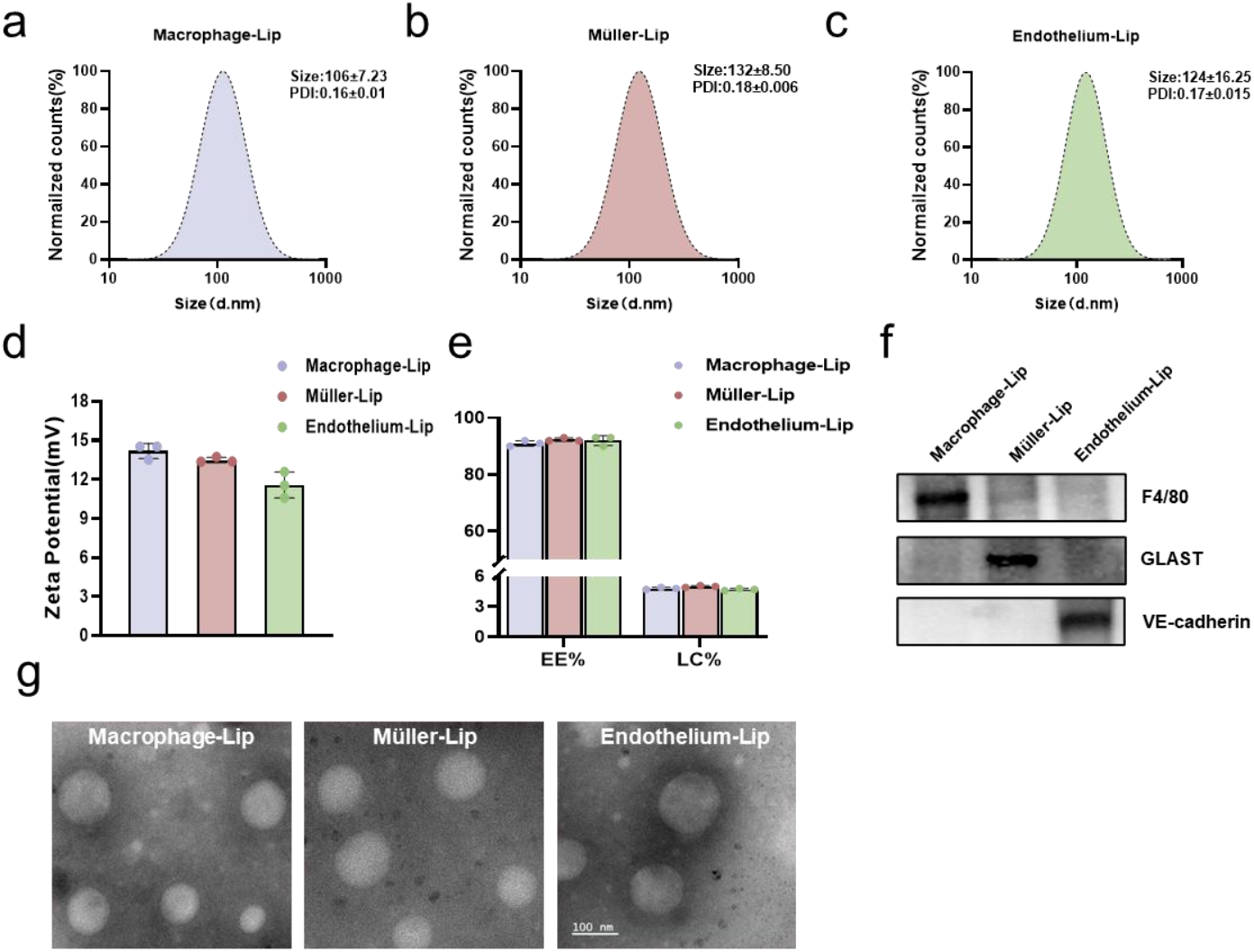
(a-c) shows the particle size distribution: (a) Macrophage-Lip; (b) Müller-Lip; (c) Endothelium-Lip; (d) Zeta potential statistics; (e) Encapsulation efficiency and drug loading statistics of Cell membrane-Lips; (f) Western blot identification of cell membrane components; (g) TEM image, scale bar 100 nm.

The siRNA encapsulation efficiency was measured using the Red I Plate™ 96 Ribo Green® RNA Quantitation Kit. As shown in Figure 1(e), the calculated encapsulation efficiencies of Macrophage-Lip, Müller-Lip, and Endothelium-Lip for siRNA were all above 92%, 93%, and 90%, respectively. The drug loading capacities were also comparable, with values of 4.8%, 5.0%, and 4.6%, respectively.

The identification of cell membrane components fused on the surface of three types of liposomes was performed by Western blot, using F4/80, GLAST and VE-cadherin as markers for macrophages, Müller cells, and endothelial cells, respectively. As shown in Figure 1(f), the presence of corresponding markers was detected in each liposome type, confirming successful fusion between cell membranes and liposomes during synthesis. The morphology of the liposomes was characterized by transmission electron microscopy (TEM), as shown in Figure 1(g), revealing uniformly spherical shapes.

### 2. Biocompatibility

Before evaluating the ocular drug delivery efficiency of the aforementioned Cell membrane-Lips, the cytotoxic effects on cells were assessed using the CCK-8 method. In this experiment, bEnd.3 cells were selected for subsequent cytological studies. Different concentrations were co-incubated with bEnd.3 cells for 24 hours. The cytotoxicity results are shown in Figure 4(a-c). The three types of Cell membrane-Lips—Macrophage-Lip, Müller-Lip, and Endothelium-Lip—exhibited no significant cytotoxic effects even at concentrations as high as 1000 μg/mL.

In vivo biocompatibility tests were performed using adult mice. After a single intravitreal injection of Macrophage-Lip, Müller-Lip, and Endothelium-Lip respectively, the mice eyeballs were enucleated 7 days later, and the retinas were isolated for pathological sectioning. The results are shown in Figure 2(d), where the retinal morphological structures remained intact in all groups without significant differences compared to the control group, indicating that all liposome formulations exhibited good ocular biocompatibility ∘

**Figure 2.**
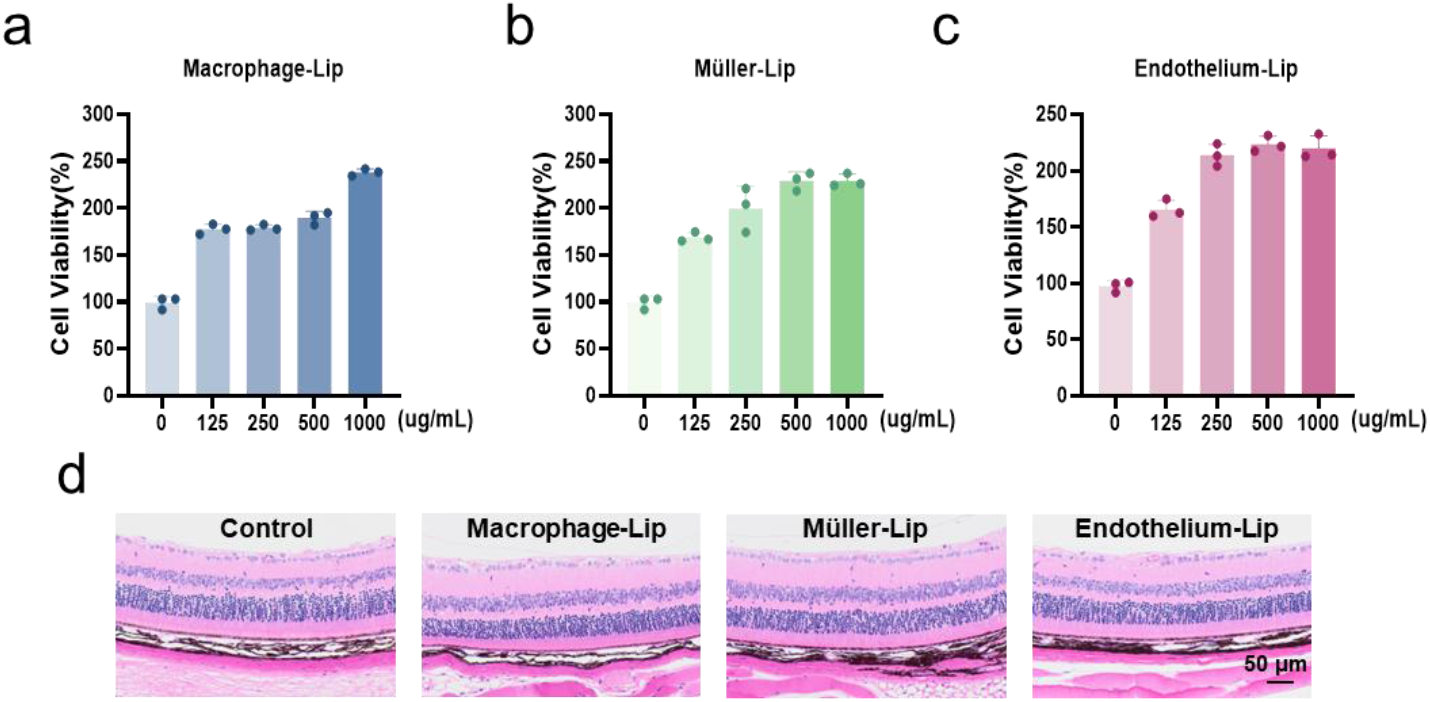
(a-c) CCK8 assay for cytotoxicity detection: (a) Macrophage-Lip; (b) Müller-Lip; (c) Endothelium-Lip; (d) HE staining of retinal tissue, scale bar = 50 μm.

### 3. Cell-uptake and Lysosomal Escape Experiments

Based on the cytotoxicity results, we selected a concentration of 150 μg/mL for subsequent validation of cellular uptake efficiency. bEnd.3 cells were treated with

Macrophage-Lip, Müller-Lip, and Endothelium-Lip for 24 hours to assess cellular internalization. As shown in Figure 3(a), after 24 hours of co-incubation, all three types of liposomes exhibited uniformly distributed red fluorescence within the cells, indicating that uptake saturation had been achieved. However, due to differences in the cell membrane components on the liposome surfaces, as illustrated in Figure 3(b), when the incubation time was reduced to 12 hours, the cellular uptake of Macrophage-Lip and Müller-Lip was significantly lower than that of Endothelium-Lip. This discrepancy in uptake may be attributed to the presence of endothelial cell membrane components in Endothelium-Lip, which likely contributed to this differential uptake behavior.

**Figure 3.**
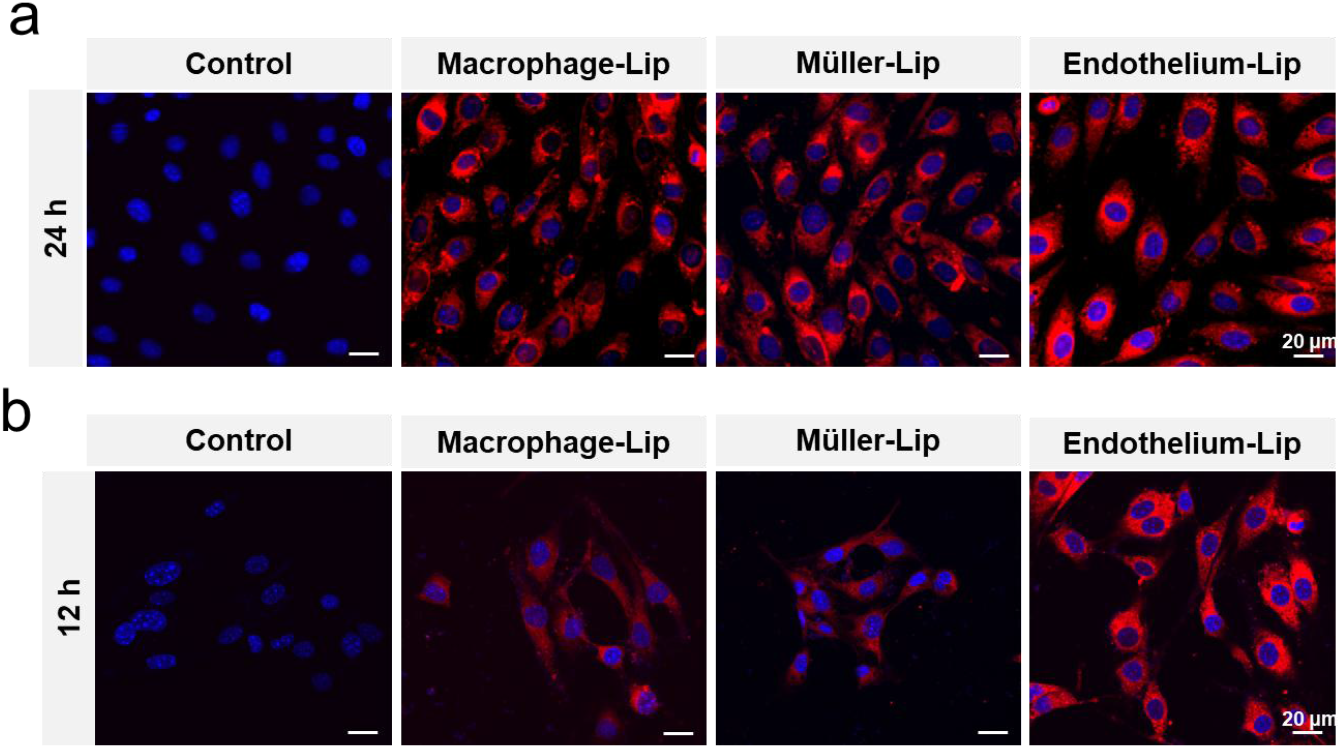
(a) Uptake results in bEnd.3 cells after 24 hours; (b) Uptake results in bEnd.3 cells after 12 hours; Liposome (red), Nucleus (blue); Scale bar: 20 μm.

Liposomes are typically taken up by lysosomes after entering cells. Based on these cellular uptake characteristics, we selected Endothelium-Lip for lysosomal escape experiments. The ionizable lipid ALC-0315, when entering lysosomes, absorbs many protons, creating a transmembrane potential difference. To achieve charge balance and concentration equilibrium, chloride ions and water from the cytoplasm flow massively into the lysosome, ultimately causing endosomal swelling and rupture - triggering the proton sponge effect ^15^. In this experiment, Lyso Tracker™ Green DND-26 was used to label lysosomes, and the colocalization of red fluorescent Endothelium-Lip with lysosomes was observed to evaluate its potential for lysosomal escape. We utilized bEnd.3 cells to validate the lysosomal escape capability of the nanoparticles. The experimental results are shown in Figure 4(a), where the colocalization phenomenon between Endothelium-Lip and lysosomes gradually decreased with prolonged incubation time. The statistical analysis of Manders’ Coefficient for colocalization is presented in Figure 4(b), demonstrating a significant reduction in the colocalization coefficient at 24 hours compared to 12 hours, with statistically significant differences. These results indicate that Endothelium-Lip is initially captured by lysosomes after cellular uptake. However, with extended incubation, the proton sponge effect mediated by ionizable lipids facilitates its escape from lysosomes.

**Figure 4.**
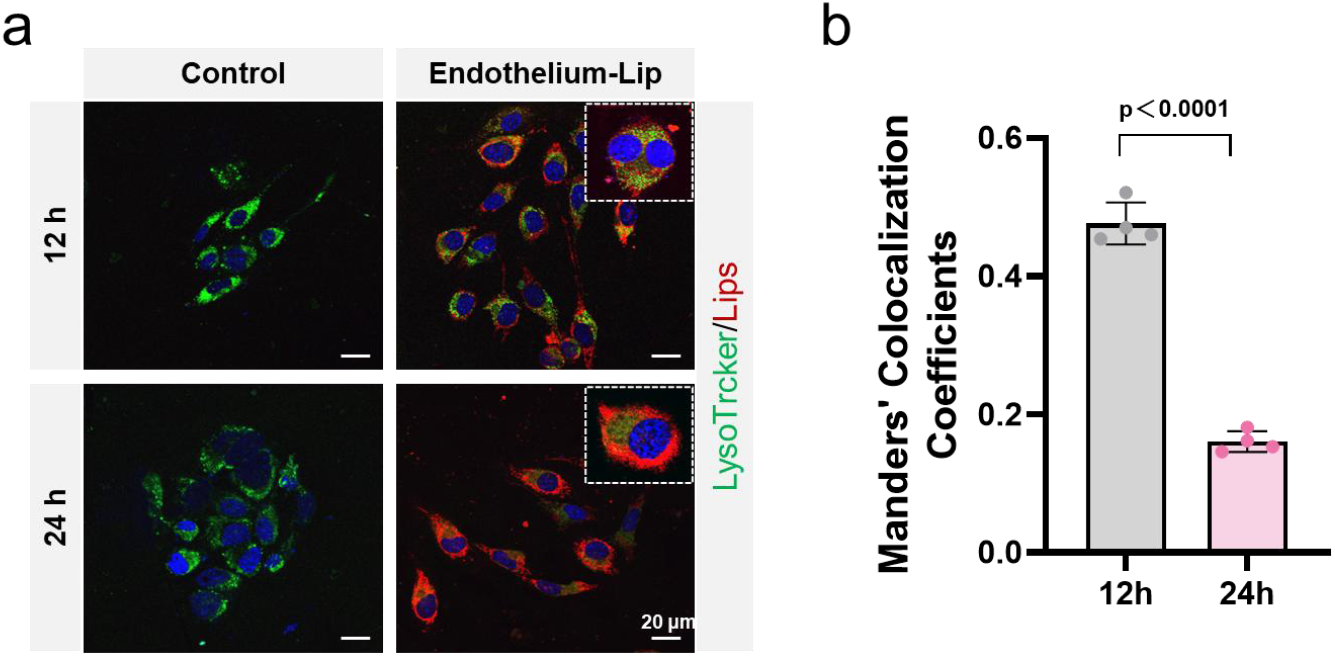
(a) Colocalization results of Endothelium-Lip with lysosomes at different time points in bEnd.3 cells; (b) Statistical analysis of Manders’ correlation coefficient. Lysosomes (green), nucleus (blue), Endothelium-Lip (red); Scale bar: 20 μm.

### 4. Targeted detection of the neovascular lesion

To evaluate the targeting efficacy of different liposomal formulations to ocular neovascularization, a laser-induced mouse CNV model was established following previously described methods. Immediately after laser induction, 1 μL each of si NC-cy3, Macrophage-Lip, Müller-Lip, and Endothelium-Lip was intravitreally injected.

Seven days later, the choroid-sclera complex was harvested for flat-mount staining, with IB4 used to label the CNV. The results are shown in Figure 5(a), where siNC-cy3 showed minimal distribution throughout the choroid. While Macrophage-Lip and Müller-Lip could penetrate the retina and accumulate extensively in the choroid, they were predominantly dispersed in the peripheral regions. In contrast, Endothelium-Lip not only efficiently traversed the retina to reach the choroid but also exhibited concentrated distribution specifically at the neovascular sites. Figure 5(b) presents a statistical analysis of fluorescence intensity at CNV sites for si NC-Cy3 and various liposomal formulations. The Endothelium-Lip group exhibited the strongest fluorescence intensity at CNV locations, with statistically significant differences compared to all other groups, demonstrating its optimal CNV targeting capability.

**Figure 5.**
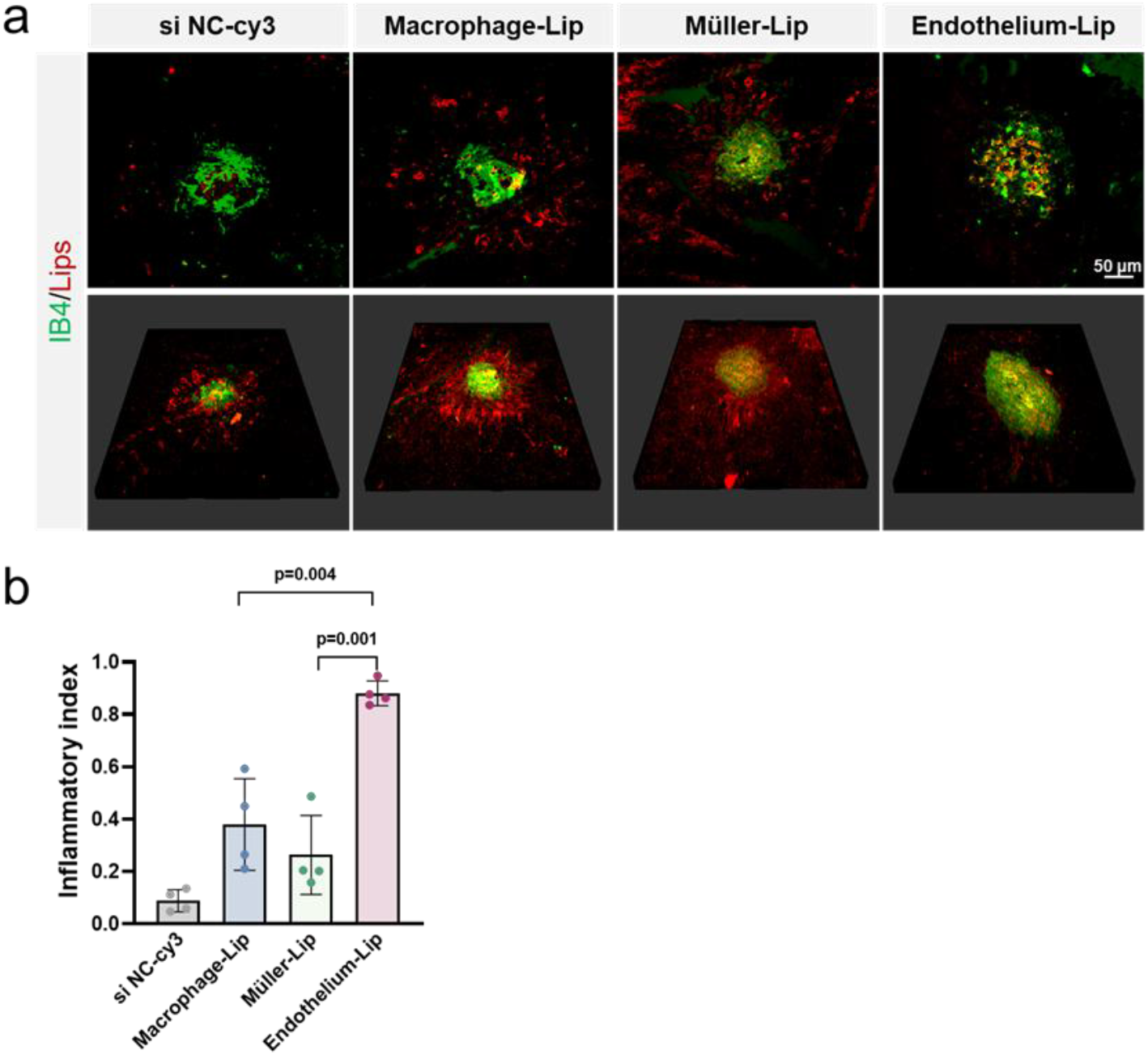
(a) Fluorescence staining images showing the distribution of liposomes in the choroidal flat-mounts 7 days after intravitreal injection, with neovascular lesions (IB4, green) and liposomes (Cy3, red). (b) Statistical results of fluorescence intensity for each group.

## Discussion

Poor response to anti-VEGF therapy remains a major challenge in the treatment and research of intraocular neovascular diseases. Although target studies have identified potential novel targets beyond the VEGF pathway, including Jagged1 in the Notch signaling pathway^16^, multiple subtypes of the Vascular Endothelial Growth Factor Receptor family^17^, and oxidative stress microenvironment^18^, their translational applications still face multiple challenges. Firstly, targets beyond the VEGF pathway (such as Angiopoietin -2 )often interact with multiple signaling networks, making single-target approaches potentially limited in efficacy ^19^. Secondly, due to the presence of the blood-retinal barrier (BRB), traditional systemic administration methods as well as non-invasive local drug delivery approaches are suboptimal for retinal tissue^20^. Intravitreal injection can directly deliver drugs to the posterior segment of the eye; however, it is also limited by injection volume, patient compliance, and the number and frequency of dosing regimens ^21^. Furthermore, neovascularization often lies in close proximity to various retinal cell types, necessitating precise targeting of pathological vessels during treatment to minimize off-target effects on surrounding healthy retinal cells ^22^.

RNA-based therapeutics offer a promising solution. Utilizing RNA drugs such as siRNA, precise silencing of multiple pro-angiogenic factors can be achieved through sequence-specific design, enabling coordinated regulation of multiple pathways^23^. Furthermore, by optimizing cell-specific delivery systems, this approach holds potential for achieving high drug accumulation at the site of pathological vessels, enhancing efficacy while reducing side effects ^24^. The main innovation of this work is the construction of a lipid nanoparticle suitable for intravitreal siRNA delivery, which exhibits excellent lesion-targeting capability and siRNA delivery efficiency.

Liposomes represent the most promising non-viral vectors. The success of Coronavirus Disease 2019 mRNA vaccines developed by Pfizer and Moderna has demonstrated their efficacy ^25^. The delivery efficiency of liposomes as carriers for RNA delivery to retinal and choroidal tissues is influenced by multiple factors including administration routes, size, charge, stability, and surface modifications. The liposomes selected for this study are composed of ionizable lipids, phospholipids, and cholesterol. ALC–0315 exhibits transient positive charge at low pH, enabling improved encapsulation efficiency with low cytotoxicity ^26^. An increase in the unsaturation of hydrophobic lipid tails from 0 to 2 cis double bonds enhances nucleic acid encapsulation efficiency ^27^. However, most LNPs primarily target the liver, while exhibiting low delivery efficiency to intraocular tissues (such as the retina)^28^. Although some studies have found that certain ionizable lipids (such as siloxane-containing lipids) can unexpectedly enhance retinal delivery, systemic delivery still faces the problem of non-specific distribution ^29^.

Based on the high water content and negatively charged environment of the vitreous cavity ^30^, this study further utilizes the weakened positive charge effect of Cell membrane-Lips to facilitate penetration through the vitreous cavity. Meanwhile, due to the existence of the “homing effect”^31^,the incorporation of cell membrane components theoretically enhances the uptake efficiency by target cells. Intraocular neovascularization is driven by localized ischemia and inflammation, involving complex pathophysiology with the participation of multiple cell types, and the neovascular components are heterogeneous^32^. Therefore, it is necessary to design and compare cell membrane types to better achieve penetration and targeting. This study selected three cell types that are both most potent and abundantly present in neovascular lesions, namely macrophages, Müller cells, and endothelial cells. Macrophages, as immune cells, possess lesion-homing properties ^33^; Müller cells, serving as the scaffold cells of the retina spanning its entire thickness, can transport ingested liposomes to the choroid ^34^; while endothelial cells act as the primary effector cells for ocular neovascularization, potentially achieving targeted delivery through membrane fusion with neovascular endothelial cells ^35^

Based on experimental results, compared to direct intravitreal injection of siRNA, all three types of Cell membrane-Lips demonstrated superior barrier penetration and lesion accumulation effects. Among them, the endothelial cell membrane-coated liposomes showed the most significant colocalization with IB4, exhibiting the best lesion-targeting capability and enabling the most efficient delivery of siRNA to CNV sites.

Although our Cell membrane-Lips demonstrate efficient targeted delivery of nucleic acid drugs to the retina and choroid, further optimization and development are still required. Primarily, it is essential to ensure the long-term safety and biocompatibility of Cell membrane-Lips, while also achieving highly efficient targeted delivery to specific retinal tissues or cells. Using scientific methods to screen different formulation compositions may yield more effective Cell membrane-Lips for retinal payload delivery. Recent studies have gradually revealed the alterations in endothelial cell surface receptors and energy metabolism under pathological conditions^36, 37^. Therefore, subsequent surface modifications or metabolic-responsive designs of Cell membrane-Lips may further enhance precise targeting capability to pathological sites.

## Conclusion

Based on the above findings, this study prepared Cell membrane-Lips through the fusion of cell membranes with liposomes. The Cell membrane-Lips possess nanoscale dimensions, a positively charged surface, and can efficiently encapsulate siRNA while protecting it from degradation by endogenous RNases. Both in vitro and in vivo settings, Cell membrane-Lips demonstrate excellent biocompatibility, rapid endocytosis, and subsequent lysosomal escape. In the mouse CNV model, intravitreal injection demonstrated superior blood-retinal barrier penetration compared to free siRNA, exhibiting enhanced lesion-targeting capabilities. Therefore, we conclude that hybrid cell membrane-coated liposomes hold great potential as drug carriers for treating ocular neovascular diseases.

## Acknowledgements

This work was supported by Key Research and Development Program of Shaanxi Province(2024SF-GJHX-38), Natural Science Foundation of Shaanxi Province (2024JC-YBMS-667), Science and Technology Program of Xi’an(25YXYJZD00018), and Subject Promotion Program of the First Affiliated Hospital of The Fourth Military Medical University (XJZT25CX52, XJZT24QN53).

## Notes

### Competing Interest Statement

The authors have declared no competing interest.

